# Pan-cancer classification of single cells in the tumour microenvironment

**DOI:** 10.1101/2022.06.14.496107

**Authors:** Ido Nofech-Mozes, David Soave, Philip Awadalla, Sagi Abelson

**Author notes:** Correspondence to &. These authors contributed equally (Philip Awadalla, Sagi Abelson).

## Abstract

Single-cell RNA sequencing reveals valuable insights into cellular heterogeneity within tumour microenvironments (TMEs), paving the way for a deep understanding of cellular mechanisms contributing to cancer. However, high heterogeneity among the same cancer types and low transcriptomic variation in immune cell subsets present challenges for accurate, high-resolution confirmation of cells’ identities. Here we present scATOMIC; a modular annotation tool for malignant and non-malignant cells. We trained scATOMIC on >250,000 cancer, immune, and stromal cells defining a pan-cancer reference across 19 common cancer types and employed a novel hierarchical approach, outperforming current classification methods. We extensively confirmed scATOMIC’s accuracy on 198 tumour biopsies and 54 blood samples encompassing >420,000 cancer and a variety of TME cells. Lastly, we demonstrate scATOMIC’s practical significance to accurately subset breast cancers into clinically relevant subtypes and predict tumours’ primary origin across metastatic cancers. Our approach represents a broadly applicable strategy to analyze multicellular cancer TMEs.

## Main

Tumour microenvironments (TMEs) are highly complex. Various immune and stromal cells within the TME interact with cancer cells to regulate processes such as angiogenesis, tumour proliferation, invasion, and metastasis, as well as mediate mechanisms of therapeutic resistance^1–4^. Single-cell RNA sequencing (scRNA-seq) techniques are explicitly suitable to disentangle complex systems as they provide transcriptome information for every cell within a sample, enabling the study of subtle transcriptomic changes reflecting different cell types and their functional states^5^.

Cell-type annotation is arguably the most critical step to derive biological insight from scRNA-seq experiments and can be performed either manually or using automatic classifiers^6,7^. Manual annotation is often an unfavorable approach as it is subjective to user definition of non-parametric clustering of cells, it’s conducted under the assumption that all the cells within a defined cluster are identical and depends on pre-existing knowledge of canonical genes. Although expression of canonical markers has been used to characterize some cell types, definitive cell markers are not always available^8^. Moreover, due to their relatively low number and the possibility of incomplete detection due to technical variation, the sole use of canonical gene expression is not ideal.

Given these limitations, there has been a recent shift towards automatic methods for cell classification, with over 100 classifiers described in a recent census of available scRNA tools^9^. To date, most existing automated annotators are focused on classification of blood or subsets of cells from other specialized tissues, thus having very limited capabilities in deciphering complex TMEs across diverse human cancers. Indeed, using single cell transcriptomics to predict cancer types and differentiate between cancer and related normal tissue cells while also classifying the large number of immune cells and stroma, is not a straightforward task^10^. In the context of TMEs, cell type predictions are challenged by high interpatient tumour cell heterogeneity among cancers of the same tissue and low transcriptomic variation among related, yet different specialized immune cells. Since cancer samples tend to cluster by patient^11,12^ and transcriptional variation is often driven by genomic instability, existing cell type classification methods which rely on distance correlations to a reference^13–15^ are expected to fail.

The current standard for the identification of malignant cells in scRNA-seq data relies on copy number variation (CNV) inference methods^16,17^. Nevertheless, these methods are incapable of providing definitive information concerning the cancer’s tissue of origin. Furthermore, CNV inference necessitates the presence of genetically unstable cells, and its accuracy may suffer when lacking a sizeable, distinctive normal cell reference within the sequenced specimen. Solely relying on the presence of inferred CNVs to annotate malignant cells may lead to false negatives or undefined cells in cancers with minimal genomic structural variation or nearly diploid genomes. Thus, a limitation in scRNA-seq analysis of tumour ecosystems is that there is no universal method for effective, detailed classification of heterogenous non-malignant TME cell types and subtypes, and cancer cells.

In this study, we present **<u>s</u>**ingle **<u>c</u>**ell **a**nnotation of **<u>t</u>**um**<u>o</u>**ur **<u>m</u>**icroenvironments **i**n pan-**<u>c</u>**ancer settings (scATOMIC), a comprehensive, pan cancer, TME cell type classifier. scATOMIC is a fully-automated, pan-cancer classification scheme that can easily be updated to capture additional subsets of normal cells and clinically relevant molecular subtypes of cancer. We devise a novel scheme that uses hierarchically organized models and elimination processes, reducing the transcriptomic complexity of the TME multi-cellular system to improve cell classification. scATOMIC will be a critical tool towards a better understanding of cancer ontogenies and the molecular interaction of diverse tumour tissues with their microenvironments.

## Results

### An overview of scATOMIC

We postulated that the sheer number of publicly available single cell transcriptomic datasets will enable the development of a highly accurate and thorough classifier for cancer, blood, and stromal cells. To define a pan cancer reference, we interrogated two comprehensive data sources containing transcriptomic-independent confirmation of cell identities. These include scRNA-seq of cancer cell lines representing 19 common cancer types^18^ and a CITE-seq dataset (proteomics and transcriptomics) of diverse peripheral blood cells^14^. scRNA-seq of stromal cells was gathered from several normal tissue sources^19–23^ (Supplementary Table 1, 2). Overall, 252,838 cells were included in the training reference dataset of scATOMIC.

Obtaining an accurate set of differentiating features is critical to successful classification, yet related cells that are functionally distinct are fated to share transcriptomic signatures (Supplementary Fig. 1). In malignant cells, neither unsupervised examination of global transcriptomic differences nor established cancer-related transcriptional programs were efficient in grouping cancer by their types (Supplementary Fig. 2), suggesting inter-patient variability as a major obstacle for accurate classification. To improve cell identity predictions, we developed a novel method, termed *reversed hierarchical classification and repetitive elimination of parental nodes* (RHC-REP) which reduces the breadth of cell types in an ensemble of classification tasks. As compared with top-down local hierarchical methods, here, predictions of terminal classes are repeatedly being evaluated to infer the cells’ broader parental nodes. During this process terminal cell classes are investigated iteratively using multiple sets of refined differentiating features until confident terminal annotations are achieved.

To develop this approach, we structured a pan cancer TME cellular hierarchy where each parent node represents a group of related cells, and each terminal node represents a single cell class of interest. Overall, we trained 21 random forest models corresponding to the total number of parent nodes (Fig. 1a). For every model, we selected differentially expressed genes (DEG) that distinguish each cell type from all other terminal classes nested within the same parent. RHC-REP prioritizes the features with the highest specificity to the interrogated cell types based on absolute expression (Methods, Fig. 1b).

**Fig. 1.**
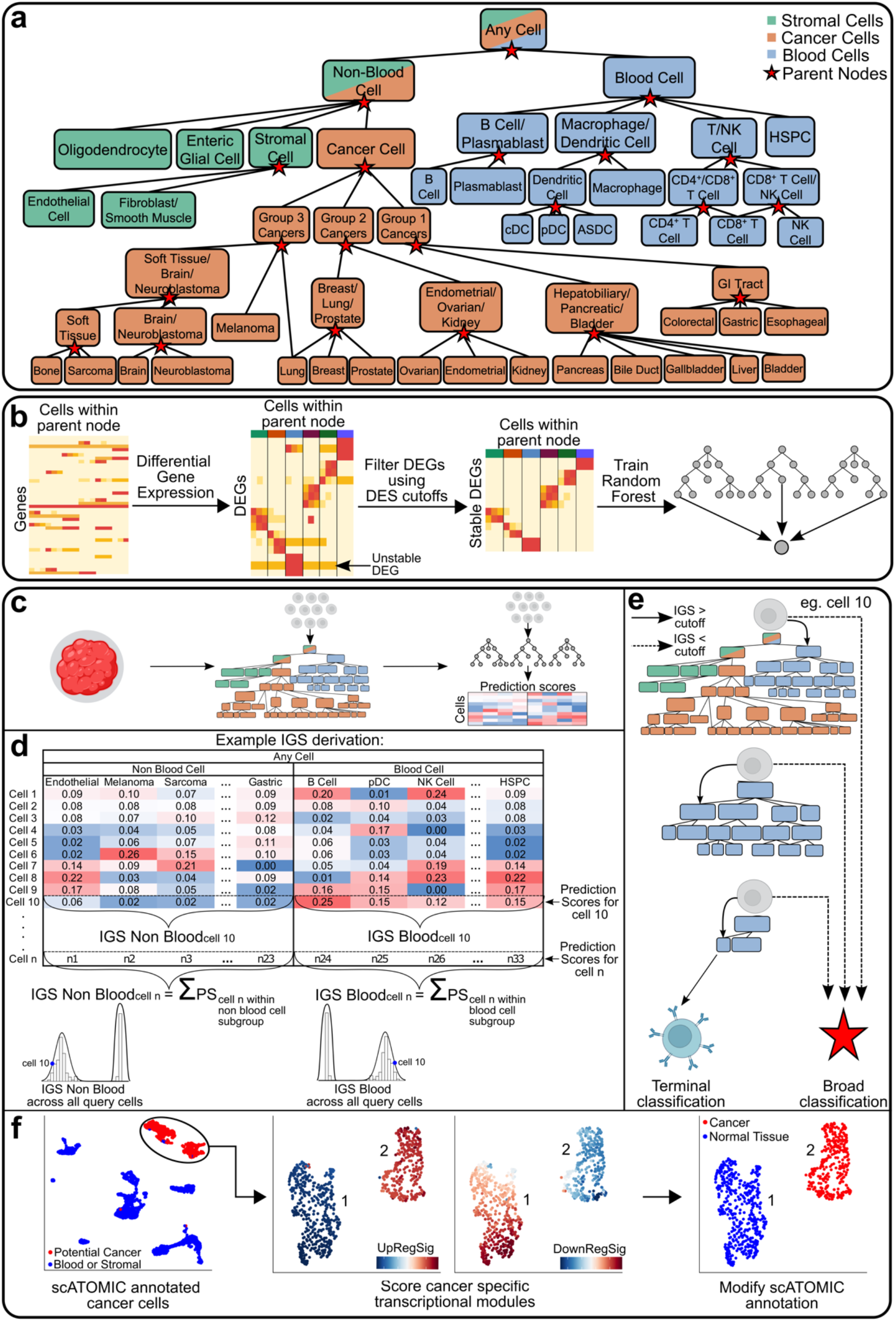
Overview of scATOMIC training and classification. **a**, Hierarchical structure of the pan cancer tumour microenvironment. The cellular hierarchies in the pan cancer tumour microenvironment are organized into a flow chart with increasing cell type resolution. Parent nodes represent broad classification branches, and terminal nodes represent specialized cell classes of interest. **b**, Training of classification branches for each parent node (n=21). The reference datasets are filtered based on transcriptomic-independent information to only include terminal cell types that are found within a particular parental node. Genes that significantly differentiate one cell type from all the others are gathered. DEGs with greater specificity to each terminal class are kept. A random forest classifier is trained on filtered, library size normalized count matrices to derive a model that provides prediction scores corresponding to the proportion of trees voting for each terminal class within the parental node. **c-f**, Classification of query datasets. **c**, Gene expression count matrices from query tumour biopsies are inputted into the first scATOMIC classification branch model, outputting a cell-by-prediction scores matrix. **d**, Prediction scores from all blood and non-blood cell subtypes are respectively summed to derive IGS distributions associating single cells with their appropriate parental class. **e**, Cells are iteratively interrogated at their next parent nodes’ corresponding models until terminal classification are obtained. Broad classifications occur if the IGS for a cell is lower than the confidence cut-off. In this example, cell 10 is subclassified until a terminal B cell designation is derived. **f**, Differentiating between cancer and tissue-specific non-malignant cells through scoring of bulk RNA-seq derived differentiating gene expression programs. scATOMIC automatically annotates population 2 as cancer cells, and population 1 as non-malignant.

During each classification task, every cell receives a vector of prediction scores (PS) corresponding to the percentage of trees voting for each terminal class in the parent node (Fig. 1c). This cell by PS matrix is then used to calculate intermediate group scores (IGS), to subsequently link cells to their next parental node in the hierarchy (Fig. 1d, Supplementary Fig. 3). At each classification task, the distribution of IGSs obtained from all the cells interrogated in the model is used to automatically define prediction cut-offs (Supplementary Fig. 4). Each cell is then interrogated by its next associated model, defined by a more discriminative set of features and fewer potential terminal classes (Fig. 1e). Cells that do not pass the IGS thresholds are given their previous parent classification and withheld from further subclassification.

Given that non-malignant cells that share the cancer’s tissue of origin can be found in cancer biospecimens (for example, normal alveolar cells in a lung biopsy) we embedded a cancer signature scoring and cell differentiating module in scATOMIC. Using an established transcriptional program scoring method^11^, cancer-type-specific up and down-regulated programs^24^ are evaluated in cells receiving an original annotation of a cancer type by scATOMIC (Methods, Fig. 1f).

### Performance evaluation and validation across internal and external datasets

To evaluate scATOMIC’s performance, we first conducted five-fold cross validation using the training reference dataset (Supplementary Table 1). We bypassed the use of IGS confidence cut-offs to ensure terminal classifications. scATOMIC achieved median F1 scores ranging from 0.91 – 0.99 across all the tested cell types (Fig. 2a), implying great accuracy in classifying the breadth of cells in the settings of pan cancer TMEs.

**Fig. 2.**
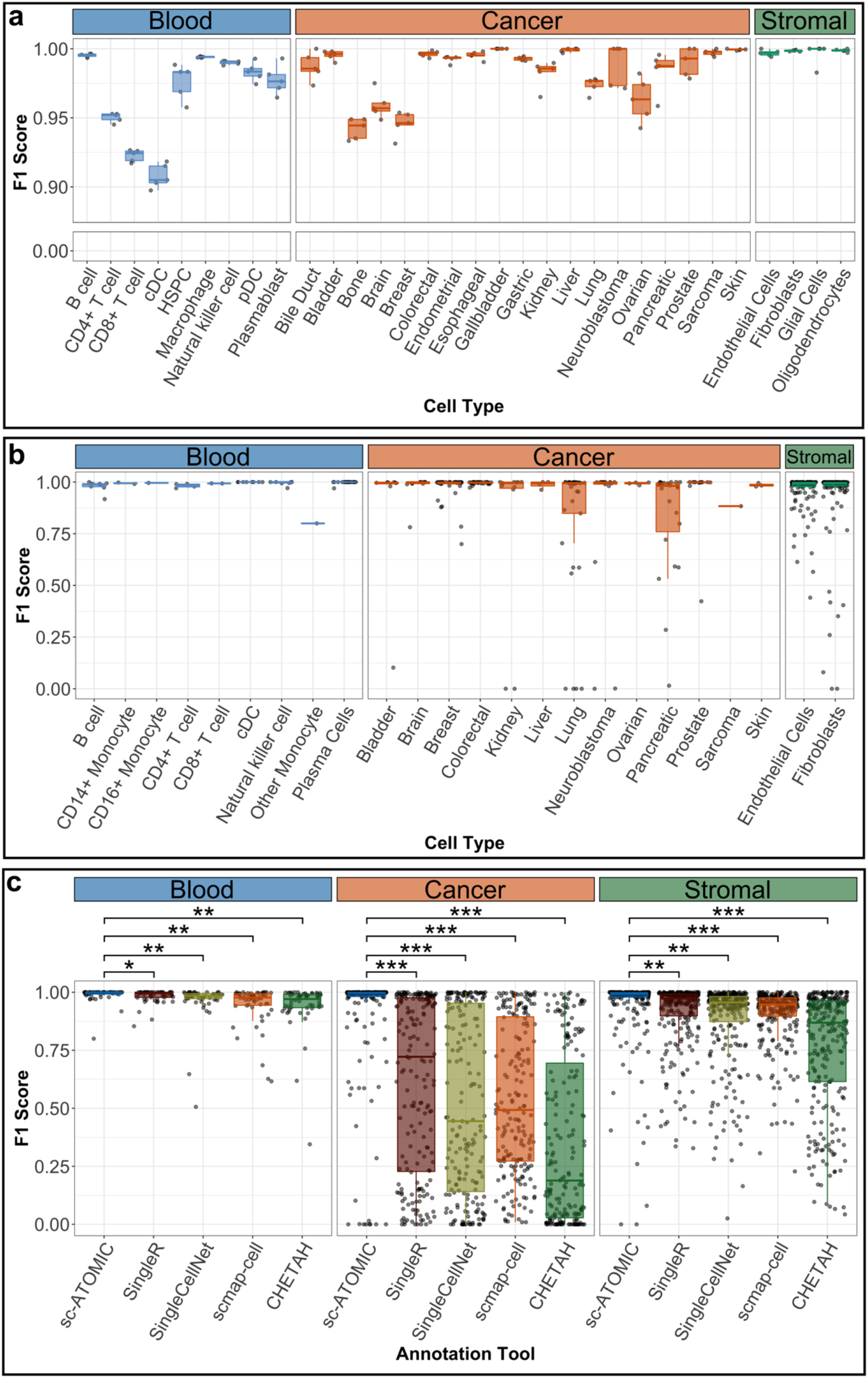
scATOMIC performs accurately in internal and external validation experiments. **a**, *k-*fold cross validation. The reference dataset was randomly split into *k*=5 sub-samples containing equal numbers of each cell type. F1 scores are shown for each cell type in each of the 5 replicates (jitter points). Each fold contained overall 50,300 cells. **b**, External validation in independent datasets. scATOMIC was validated in scRNA-seq datasets of flow cytometry sorted blood cells, and aneuploid cancer cells and stromal cells from primary tumour biopsies. F1 scores are shown for each cell type within individual samples (jitter points). A total of 424,534 cells were used. **c**, scATOMIC outperforms other existing automatic cell type annotators, particularly when applied to identify cancer cells and determine their type. Four existing classifiers were provided the same training/reference and external validation datasets as scATOMIC. Combined F1 scores for each of the three major cell class, blood, cancer and stroma are shown (jitter points). (* P < 0.05, ** P < 1.1 × 10^−6^, *** P < 2 × 10^−16^, Wilcoxon Rank Sum Test). For all plots, boxes and whiskers represent the lower fence, first quartile (Q1), median (Q2), third quartile (Q3), and upper fence.

We next aimed to conduct a comprehensive external validation of scATOMIC performance. To build an independent validation dataset with high confidence cell annotations, we mined publicly available scRNA-seq data from primary tumour biopsies and blood samples. Overall, the curated set used for validation contained 228,296 cancer, 81,965 stroma and 114,273 blood cells from 198 primary biopsies spanning 13 cancer types, and 54 blood samples (Supplementary Table 3). Importantly, this ground truth set includes cancer cells supported by abnormal CNV profiles, and non-malignant cells that were either selected using magnetic beads or were flow sorted. Similar to the results obtained from internal validation, in this independent validation process, scATOMIC achieved a median F1 score of 0.99 across all cell types (Fig. 2b). Overall, these results demonstrate the broad abilities of scATOMIC core algorithm to detect cancer cells and their type, as well as predicting non-malignant cell types and subtype.

### Comparison with other cell-type classification methods

We compared scATOMIC’s performance to four commonly used scRNA-seq classifiers (SingleR, SingleCellNet, scmap-cell, and CHETAH)^13,15,25,26^. These annotators, altogether, represent reference correlation, random forest, flat and hierarchical classification methods making their comparison with scATOMIC’s underlying RHC-REP approach highly suitable. Each tool was provided with the same reference and external validation dataset as scATOMIC for training and testing (Supplementary Table 1-3). All classifiers were highly accurate in classifying non-malignant cells (median F1>0.91, Fig. 2c). However, for cancer cells in particular, F1-scores were significantly lower as compared with scATOMIC. The next best performing classifier following scATOMIC was SingleR with median F1 scores of 0.96, 0.97 and 0.72, for blood, stromal, and cancer cells, respectively (Fig. 2c). Using scATOMIC, we obtained median F1 scores of 0.99 in each of these categories. As existing cell type classification tools were not designed to annotate malignant cells, this comparison highlights the novelty of scATOMIC’s unique ability to overcome the complexity presented in pan cancer settings to accurately identify cancer cells while also having significantly better performance in annotating stroma and blood.

### Distinguishing between non-malignant, tissue-specific cells and cancer

Aneuploid CNV profiles are highly associated with the development and progression of numerous cancers by impacting gene expression levels^27^. We assessed scATOMIC’s ability to distinguish malignant cells from other normal cells of the TME, sharing the same cell-of-origin, by comparing scATOMIC’s final cancer predictions (Fig. 1f) to their inferred CNV-based ploidy status^17^ (Methods). We observed strong agreement in cells predicted as malignant- and aneuploid-inferred CNV profiles across biopsies, as well as between non-malignant detected cells and diploid inferred profiles, with a median agreement rate of 86.5% (Fig. 3). Discordant cases, defined here as cases with an agreement rate below the 1^st^ quartile (Q1 ≤ 63.2%), were typically attributed to tumours with low CNV levels, intra-tumoural malignant subclones, low number of cancer cells harboring the CNV, or low number of reference cells (Supplementary Fig. 5). In only 7% of discordant cases more cells were inferred as aneuploid than cells annotated as malignant by scATOMIC (Fig. 3b). These results suggest that cancer and related normal tissue cells can efficiently be classified by their transcriptomic profiles using scRNA-seq data, independent of their ploidy status.

**Fig. 3.**
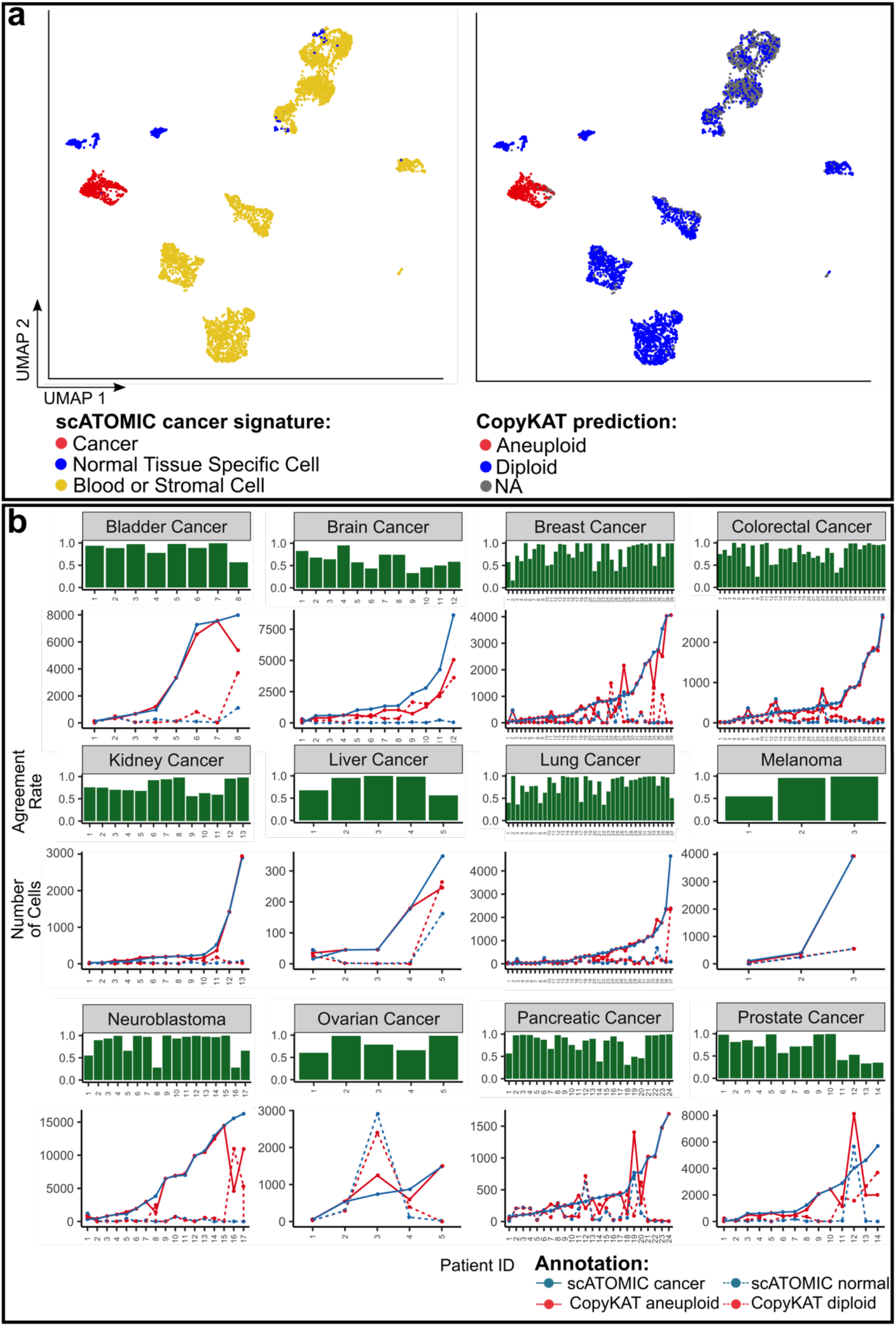
scATOMIC effectively distinguishes between malignant cells and normal tissue specific cells. **a**, scATOMIC predictions and inferred ploidy in breast cancer patient CID4066^34^. Cells are coloured by scATOMIC predictions and CNV-based inferred ploidy. A scATOMIC-predicted malignant cells are inferred as aneuploid cells while normal tissue cells are inferred as diploid. **b**, Comparison of scATOMIC cancer predictions and inferred ploidy statues across the external validation dataset. Blue points represent the number of cells predicted as malignant (solid line) and non-malignant (dashed line) by scATOMIC. Red points represent the number of cells inferred as aneuploid (solid line) and diploid (dashed line). Green bars represent agreement rate in each biopsy. Rates do not include cells without a confident ploidy status (that is received an “NA” annotation by CopyKAT). For visualization purposes only, the figure does not include the single sarcoma case (agreement rate = 93.9%)

### scATOMIC annotations increase cellular resolution in tumour biopsies

To further demonstrate the benefits of scATOMIC in annotating multi-cellular TMEs, we analyzed several datasets, including scRNA-seq of lung cancer^28^. Original annotations for this dataset were determined using SingleR^13^ with its default references and manually augmented using a combination of cell type signatures and known canonical markers (Fig. 4a). Similar to our observations (Supplementary Fig. 1) Slyper et al (2020), noted that overlapping expression programs between T cells and NK cells make these cell types difficult to accurately discriminate. scATOMIC resolved T cells to CD4+, CD8+ and NK cells as well as other challenging cell types including plasma cells, and plasmacytoid dendritic cells (pDCs) which scATOMIC separated from B cells (Fig. 4a). Unsupervised clustering and the expression of cell specific markers supported the existence of these separated cell identities (Supplementary Fig. 6). Of note, this biopsy included mast cells, a cell class which was not included in scATOMIC’s training reference. Using scATOMIC’s automatic approach to set confidence IGS cut-offs (Supplementary Fig. 4), it abstained from falsely annotating these cells, determining them as unknown. Lastly, scATOMIC resolved epithelial cells into lung cancer cells and normal tissue cells.

**Fig. 4.**
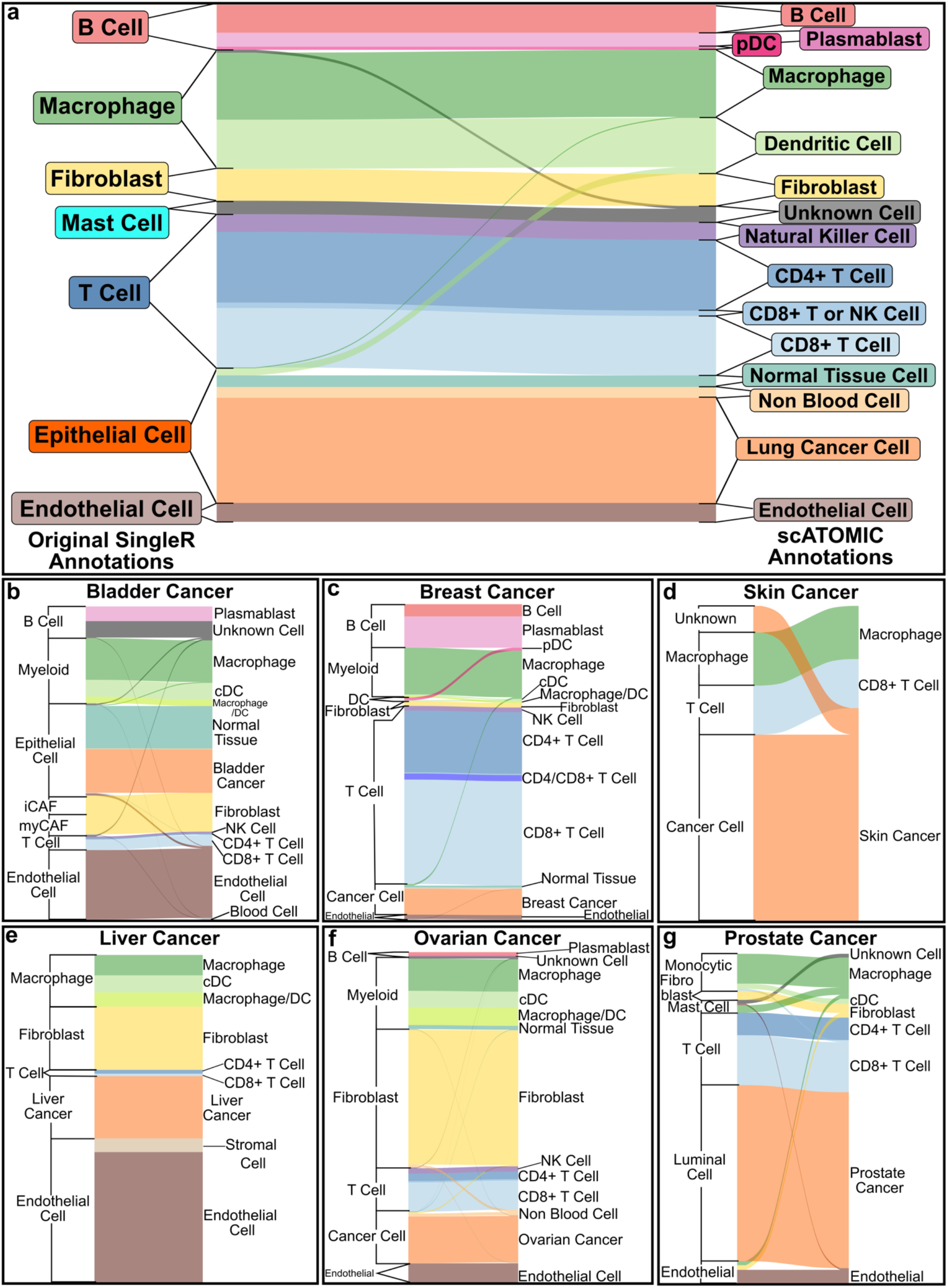
scATOMIC provides greater cellular resolution than original annotations across tumour datasets. **a**, Sankey plot comparing original cell type annotations to higher resolution scATOMIC annotations in a recent lung cancer biopsy dataset^28^. scATOMIC identifies lung cancer as the tissue of origin and distinguishes these cells from normal lung tissue cells. scATOMIC identifies subtypes of blood cells. **b-g**, scATOMIC identifies the tumour origin of common cancers and deliver relatively high resolution in other cell types^4,29–32^. Colours represent the scATOMIC annotations. The height of each block represents the relative number of cells that received a respective annotation.

Increased cellular resolution was also observed in other recent datasets of different cancer types including bladder^4^, breast^29^, liver^30^, ovarian^29^, prostate^31^, skin cancer^32^ (Fig. 4b-g). Additionally, scATOMIC identified hematopoietic stem/progenitor cells (HSPCs) in glioblastoma^33^ (Supplementary Fig. 7); a population which was shown promote tumour cell proliferation^33^.

Collectively, this analysis demonstrates the ability of scATOMIC’s core hierarchical algorithm to resolve cell identities at high resolution, identify rare cell types, abstain from falsely classifying unknown cells, and determine cancer types.

### Extending the core scATOMIC hierarchy for novel applications

By leveraging RHC-REP, one can easily deploy new scRNA-seq data to train extensions at any terminal branch of the hierarchy. We thought that extending the breast cancer classification node would provide a practical example of utilizing modularity (Fig. 5a). Two sizable scRNA-seq breast cancer atlases were used to train, and independently test (Supplementary Table 4, 5) a classification model that resolves breast cancer cells into the major ER+, HER2+ and triple negative breast cancer (TN) histological subtypes. scATOMIC correctly subtyped 39 of the 40 (97.5%) training-independent breast cancer biopsies spanning ER+, HER2+, ER+/HER2+, and TN tumours; as determined by immune-staining^34,35^ (Fig. 5b, Supplementary Table 5).

**Fig. 5.**
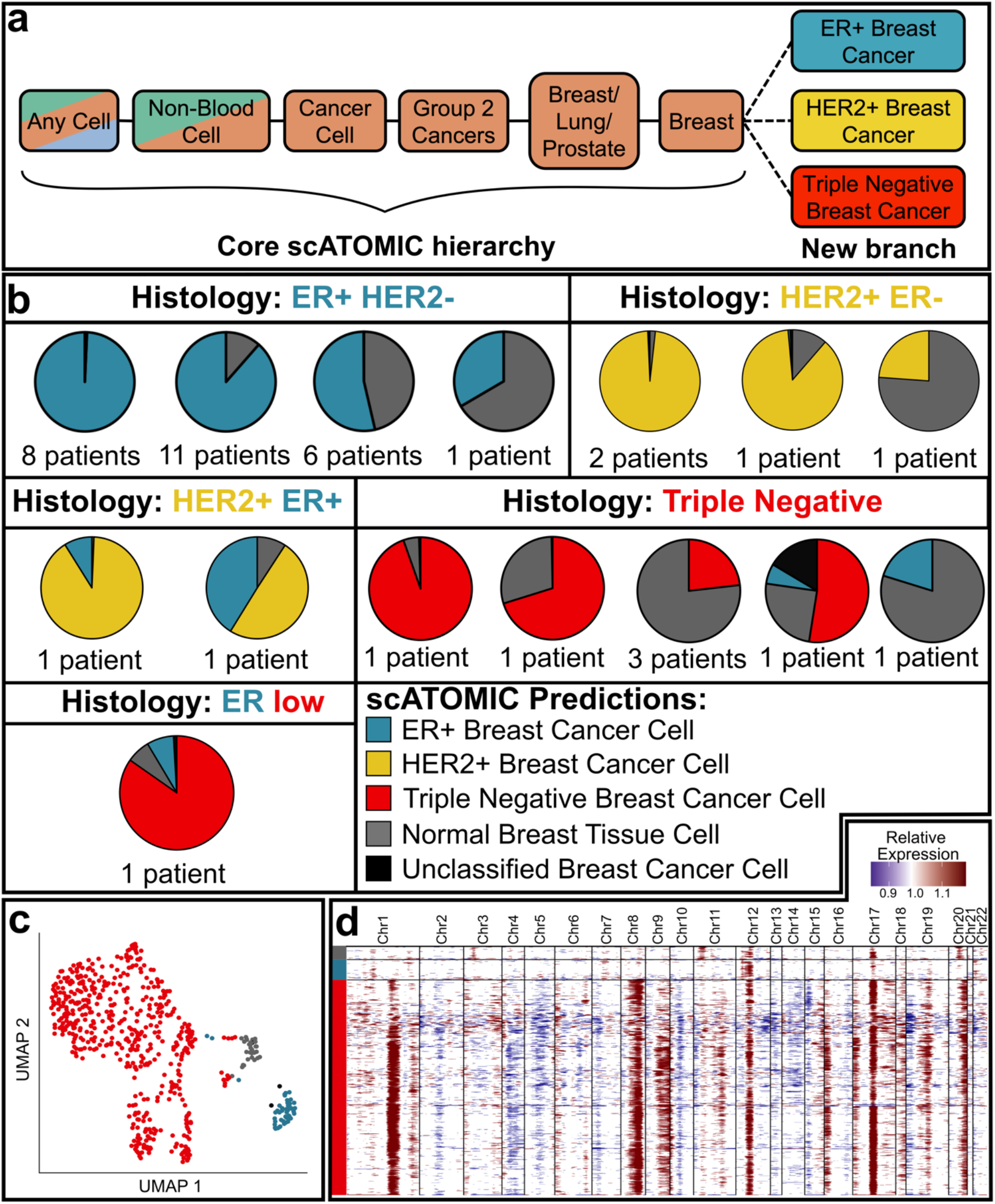
Extending the core scATOMIC model to further classify breast cancer subtypes. **a**, The terminal breast cancer cell node from the core hierarchy of scATOMIC is extended to subclassify breast cancers into their major ER+, HER2+, and triple negative histological subtypes. **b**, Validation of scATOMIC predictions in an external cohort^35^. Pie charts reflecting intratumoural breast subtype heterogeneity according to scATOMIC classification are shown for each reported histological subtype. Patient specimens with similar distributions of cell annotations are illustrated together in a single pie chart. **c**, Breast cells from an ER-low tumour (Patient: ER-AH0319) are visualized on UMAP and coloured by scATOMIC predictions. ER+ breast cancer cells represent a sub-clonal cancer cell population. **d**, Inferred CNV profiles of cells from ER-low tumour. Red represents inferred gains, while blue represents inferred losses of genomic regions. The y axis is coloured according to scATOMIC prediction. Colours representing scATOMIC predictions apply to all the panels in the figure.

We observed different degrees of tumour cellularity, with 6 biopsies (15%) having more predicted normal breast cells than cancer cells. HER2+/ER+ double positive samples received mixed annotations of HER2+ and ER+ cells demonstrating the ability to correctly identify double positive tumours. In another tumour reported as ER^low^ (that is, <10% ER+ cancer cells by immunostaining), scATOMIC identified 8% ER+ breast cancer cells (Fig. 5c, Supplementary Table 5). Of note, scATOMIC identified these ER+ cells as a malignant clone, in line with the histology report, however CNV inference predicted a diploid profile (Fig. 5d). This example highlights a distinct subpopulation of cancer cells that could have been misinterpreted as normal tissue by strictly relying on CNV inference, thus suggesting integrative approach for best results. Overall, these data demonstrate scATOMIC’s practical and modular framework to further subset primary tumour classes into their clinically relevant subtypes.

### scATOMIC identifies the tumour of origin across metastatic cancers

Given that existing single cell annotation tools are not designed to provide information regarding the originating tissue of a cancer cell, we applied scATOMIC’s unique ability to predict the tumour origin in settings where it may be unknown. We curated a dataset of 62 metastatic biopsies from breast, kidney, lung, ovarian, and skin cancers from diverse anatomical sites (Supplementary Table 6). In 54 of the 62 samples (87.1%), the primary tissue of origin was correctly predicted by scATOMIC (Fig. 6), demonstrating its robustness at distant sites, in cells that may have undergone transcriptional changes associated with metastasis. In 6 samples (9.7%), the predicted cancer type and the reported primary were related cancers falling under the same immediate parent node. For example, a mixed serous/clear-cell ovarian carcinoma was predicted to be endometrial cancer, with relatively low classification scores (Supplementary Fig. 8). Two of these 6 biopsies included rare cancer subtypes that were not represented in the reference cell lines used for training (Supplementary Table 2). One (1.6%) lung cancer was misclassified as pancreatic cancer and lastly, in one lower throughput melanoma scRNA-seq, with only 6 reported cancer cells^11^, scATOMIC found none. Overall, these results show that accurate detection of metastatic cancers’ tissue of origin using single cell transcriptomics is feasible. Furthermore, scATOMIC can aid in identifying cancers’ primary site in a variety of solid human tumours.

**Fig. 6.**
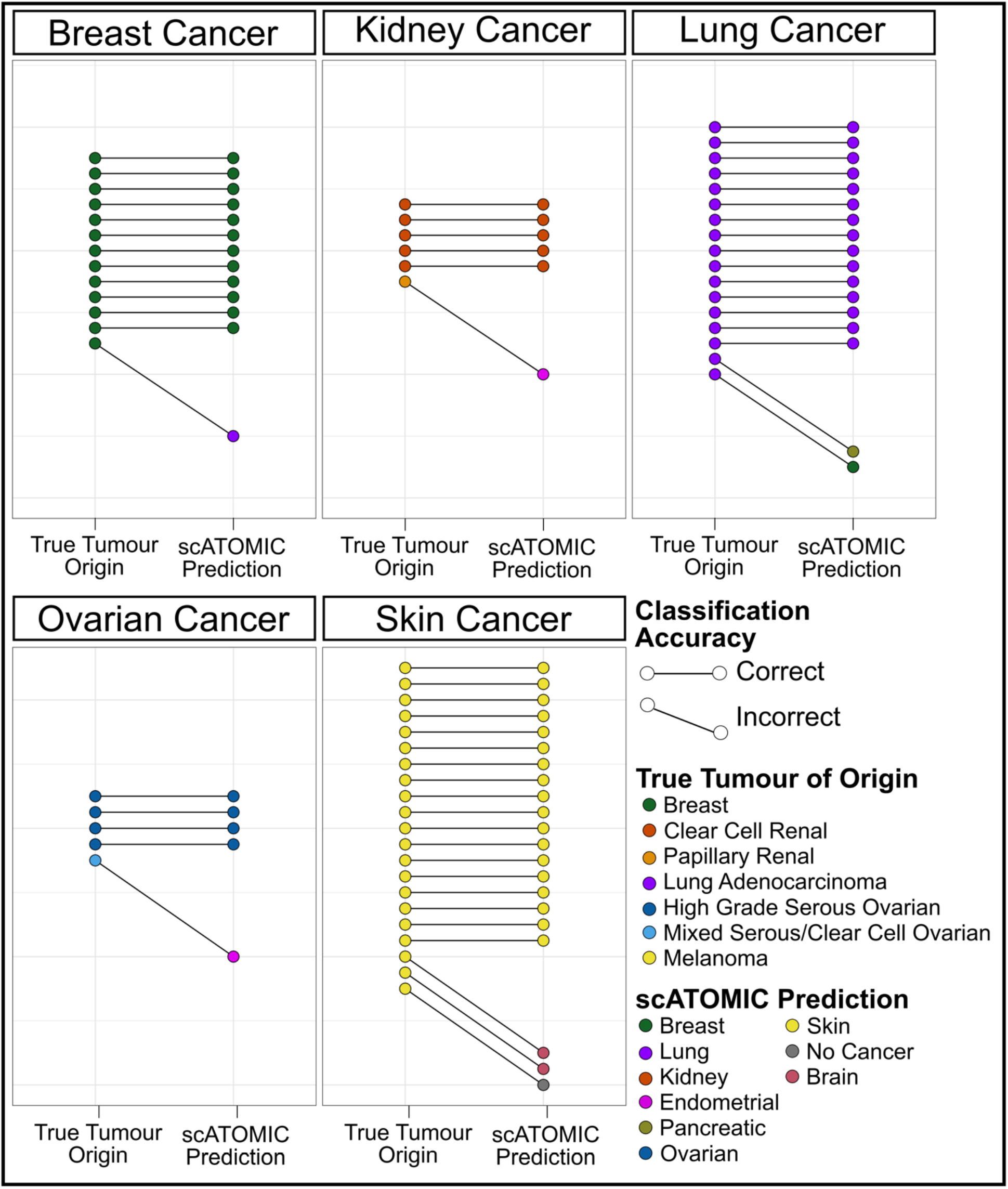
scATOMIC accurately identifies the tissue of origin in metastatic tumour biopsies. scATOMIC was applied to 62 metastatic tumours from breast, kidney, lung, ovarian and skin. Metastatic sites included the brain, lungs, GI tract, liver, adrenal glands, lymph nodes, abdomen, and peritoneal cavity. Each pair of dots represents the true tumour origin and the predicted origin. Horizontal connected lines represent correct predictions, while diagonal lines represent incorrect predictions. True tumour origins are coloured by the reported cancer subtype.

## Discussion

The rate of scRNA-seq publications reporting major scientific insights concerning the function of various immune and stromal cells in cancer has increased steadily over the years^29,36,37^. However, the lack of an automated method that can also standardize the identification of single malignant cells is becoming a major obstacle to accurately study tumour-microenvironment interactions across various cancer types.

We developed scATOMIC, a novel tool to annotate the TME in pan cancer settings, unlike other existing methods tested, scATOMIC can overcome several classification challenges, including high inter-patient heterogeneity and highly overlapping expression profiling among specificized immune cells. By using stably expressed transcripts as features, novel structured classification, and models trained using reliable and large datasets, scATOMIC has proven to be the only tool suitable for accurate identification of cancer cells and their origin. Moreover, scATOMIC outperforms other existing automatic cell type annotators when classifying blood and stromal cells using our training reference. In samples with genome instability and an appropriate reference of normal cells, we found high agreement between scATOMIC and the current leading CNV inference approach to pinpoint malignant cells in scRNA-seq data. However, in samples with cancer sub-clones defined by variable CNV burdens, cancer cells with near diploid genomic profiles, or few normal cells to serve as controls, CNV inference may fall short. Since information concerning CNV may be use for cancer prognostication, and a degree of discordancy still exists, using these two independent approaches to annotate cancer cells and their type is recommended.

We designed the core, ploidy-neutral scATOMIC algorithm to accurately identify cancer and normal tissue cells across 19 common cancer types, including key rare populations such as plasmacytoid dendritic cell and hematopoietic stem and progenitor cells that were reported in cancer tissues and are associated with immunosuppressive phenotypes^28,33^. To ensure that scATOMIC remains powerful, we designed it in a way that new data can be easily interrogated to extend the core hierarchy by adding new terminal cell classes. We demonstrated this modularity by further classifying breast cancers into their clinically relevant molecular subtypes achieving extremely high agreement between transcriptomics and immunostaining. With the progressive accumulation of high quality publicly available scRNA-seq data, future extensions of the core hierarchy to also subclassify immune cells to their more resolved states will become simple. We expect that scATOMIC’s flexible approach to train additional terminal classes will expedite broad cancer studies seeking to refine prognostication or cell–cell communication from single cell transcriptomic information.

As molecular classification of cancers by tissue-of-origin is fundamental to diagnostic pathology we demonstrated scATOMIC’s ability and high accuracy in predicting the primary origin of metastatic tumours. Additional work is required to evaluate the limits of single cell transcriptomics to predict cancer origin, specifically in a cohort of cancers of unknown primary and other contexts where distinguishing primary and metastatic tumours is not trivial, such as mucinous ovarian carcinoma^38^.

In summary, we have described, benchmarked, and validated a highly accurate single cell annotation tool across TMEs of 19 common and deadly cancer types. The novel classification approach underlying scATOMIC used here to tackle the complexity associated with multicellular TMEs might be of interest in areas other than the cancer field entailing multi-class structured systems. We highlight the unique benefits that scATOMIC holds in cancer settings compared to other tools, providing a method to standardize single cell cancer transcriptomic studies.

## Methods

### Defining a pan cancer tumour microenvironment cell type hierarchy

We defined the structure of the hierarchy and identity of parental nodes in the core scATOMIC algorithm (Fig. 1a) based on known biological relationships among cell types. In some instances, where identifying the relationship between cell types was not trivial, we interrogated which cell classes shared high PS in the first classification branch of scATOMIC using a subset of data from Slyper, M. *et al*^28^ and defined associations between cell classes. For example, similar classification scores were initially found in cancers originated in related organ systems or cancer cells that share histological subtypes (e.g., clear cell carcinoma) yet may arise in different tissues^39^. In the case of immune cells, we found that overlapping classification scores mostly recapitulated the established hematopoietic cellular hierarchy^40^.

### Feature Selection

Raw gene by cell count matrices were gathered from multiple sources (Supplementary Table 1) and were organized into 21 parent groups (Supplementary Fig. 3). To merge matrices into a particular parent dataset, we removed genes that are not represented in all of the data sources. In each parent dataset we removed cells with less than 500 expressed genes (as defined by non-zero counts) or with more than 25% of their reads being mapped to mitochondrial genes.

To find DEGs between each terminal cell type and all other terminal cell types present in the same parent node we used the ‘Seurat’ R package v4.0.1^14^. Raw gene by cell count matrices were normalized and variance stabilized using the *SCTransform* function to remove technical variability. Principle components were found using the *RunPCA* function, on the “SCT” assay. Louvain clustering was performed by first applying the *FindNeighbors* function on the top 50 PCs, followed by the *FindClusters* function with a resolution of 2. The identity of the resulted cell clusters was determined by the transcriptomic-independent ground truth associated with the training datasets. For each model, DEGs were found using the *FindMarkers* function with ident.1 set to include all clusters containing a particular terminal cell type and ident.2 being all other cells in the parent node. The function returned a list of DEGs per class that passed default Seurat filtering settings: a log_2_ fold-change of at least 0.25, and at least 10% of the cells in ident.1 or ident.2 expressing the respective gene. We defined a differential expression score as the difference of the fraction of cells expressing a non-zero value for a respective DEG in ident.1 and ident.2 (DES = pct_expr_ident.1 – pct_expr_ident.2). For each terminal cell type, we kept genes with a DES greater than the mean DES of all DEGs for that cell type. We removed all ribosomal genes. We also removed DEGs that had a pct_expr_ident.2 greater than 40% to ensure high performance when interrogating datasets with large technical variation. For the same reason, we set a minimum and maximum number of DEGs for each cell type at 50 and 200. Specifically, a minimum number of features was set to mitigate potential issues in classifying cells with high levels of technical dropout. In the case where there are fewer than 50 DEGs with DES higher than the mean, we kept the top 50 DEGs ranked by DES. Features that were used for each cell class at each classification layer are provided in Supplementary Table 7.

### Random forest training

To minimize bias associated with imbalanced classification towards majority classes^41^, we randomly sampled an equal number of cells from each terminal class, with replacement. Library size of each single cell was normalized by using the *library*.*size*.*normalize* function from the ‘Rmagic’ v2.0.3 package^42^. Prior to each model training, normalized counts were filtered to include selected features and cells within the corresponding parental node. Read count values were transformed to a fraction of the total filtered counts. A random forest classifier was trained on the transformed matrix using the ‘randomForest’ R package v4.6-14 with 500 trees and default parameters^43^. The specific cell type organization of the 21 classifiers is detailed in Supplementary Fig. 3.

### Running scATOMIC on query datasets

Raw gene by cell count matrices were filtered to remove cells with non-zero counts for less than 500 genes or with more than 25% of their reads being mapped to mitochondrial genes. We imputed missing values in DEG using the *magic* function from the ‘Rmagic’ package^42^, where all the cells within the query dataset act as a reference. In some datasets where there were no reported values for specific selected features, we assign a value of zero before imputation. Following each classification task, every cell receives a vector of prediction scores corresponding to the percentage of trees voting for each terminal class in the random forest model. The values in each vector within the next immediate parent node were then summed to generate IGSs. For example, if the first model interrogating all the terminal classes in the hierarchy is running (i.e. the parent model “Any Cell”), the output for each cell will be composed of two intermediate group scores. The first corresponds to the sum of trees voting to all the terminal cell classes belonging to the “Blood Cell” parent node and the second IGS for the “Non-Blood Cells” parent node (Fig. 1d). Data from all the cells in the interrogated sample was used to derive IGS parent distributions. Cells that receive an IGS greater than the defined parent threshold continue down the classification hierarchy until they are terminally classified (Fig. 1e). At any stage, if a cell receives an IGS lower than the calculated threshold it will be annotated based the previous parental node. We automatically determine an IGS threshold for a classification to be deemed confident (Supplementary Fig. 4). Using the *ajus* function from the ‘agrmt’ package v1.42.4^44^, we classify IGS distributions for each IGS calculated among all cells within a parental node as either unimodal or bimodal. Unimodal distributions suggest a layer includes one subtype, while bimodal distributions indicate there is likely more than one subtype. For unimodal distributions we set the IGS threshold to be the mean IGS (*µ*) – 3 standard deviations (*σ)*. For bimodal distributions, using the *em* function from the ‘Cutoff’ R package^45^ v0.1.0, we fit a mixture model for the distributions and predict estimates of mean and standard deviations for both distributions using the expectation maximation algorithm. We selected a conservative approach when a mixed cell type population exists in a layer by setting the IGS threshold in bimodal distributions to be the mean of the higher score modality (µ_2_) – 2 standard deviations (*σ*_2_). We set the maximum IGS threshold to be 0.7, as in some distributions, such as when a single pure population remained for classification, unreasonably high thresholds may be obtained.

### Scoring cancer signatures to refine cancer cell predictions and identity of normal tissue-specific cells

Lists of genes differentially expressed between different cancer types and their matched normal tissues were obtained from OncoDB^24^. We selected DEGs with a log2 fold change greater than 1 or less than -1 and an adjusted P-value less than 0.01. Since OncoDB is based on bulk RNA-seq, we further filtered the DEG list to only include those with reported expression values in the query scRNA-seq dataset. Upregulated gene programs and downregulated gene programs^24^ were scored using the AddModuleScore function from Seurat^11,14^ in each cell predicted as cancer by random forests. Ward.D2 hierarchical clustering was then performed on a Euclidean distance matrix of each cell’s upregulated and downregulated cancer programs. Two groups are derived using the cutree function. At this stage, scATOMIC evaluates in each group the percentage of normal cells, corresponding to those cells annotated as either blood, stroma or cancer cells with lower upregulated programs score compared to downregulated programs score. Since scores are relative across cells in the whole dataset, we then filtered out all normal cells in the group with a greater percentage of normal cells and repeated scoring of cancer programs, hierarchical clustering, calculating the percentage of normal cells and filtering. These steps are repeated until both clusters do not contain more than 20% normal cells. Cells that were initially classified as cancer which were scored as normal cells are given a normal tissue cell label.

### Benchmarking scATOMIC

Internal validation was performed by splitting the training dataset (Supplementary Table 1) into five subsets containing equal proportions of each cell type. For each iteration we used 4 subsets as scATOMIC training dataset and applied scATOMIC to a held-out independent test subset. scATOMIC was configured to refrained from using IGS thresholds to ensure that terminal classifications are obtained for each cell. F1 scores were calculated for each terminal cell type at each iteration (Supplementary Table 1). AXL^+^ dendritic cells (ASDC) represent a rare, recently discovered, transitional state between cDCs and pDCs^46^. Due to their small numbers in the training data, these cells were omitted from the internal validation procedure.

For external validation of scATOMIC we used 424,534 cancer, blood and stromal cells gathered from various sources (Supplementary Table 3). For ground truth, we relied on the authors’ annotations of stromal cells. To improve reliability, cells that were annotated as cancer by the authors were subjected to additional validation by CNV inference using CopyKAT^17^. For blood cells, we used flow sorted or magnetic bead enriched datasets from peripheral blood or bone marrow. We excluded cell types from individual samples if their number was less than 30. As with the internal validation, we avoided intermediate annotations by omitting IGS confidence thresholds. Per sample F1 scores were calculated for each terminal cell type.

### Comparison between scATOMIC and other tools

The same training and validation datasets provided to scATOMIC were also used to evaluate the performance of all the other tested tools. A SingleCellNet^25^ random forest model was trained on a balanced reference of 2500 random samples of each cell type, using default parameters. SingleCellNet expects a class-balanced as input. We provided the same class-balanced matrix used in scATOMIC’s first classification model, representing all the cell classes. CHETAH^26^ is using an alternative hierarchical classification approach. We used CHETAH with its default settings with the exception of the ‘thresh’ parameter that was set to zero to enforce terminal annotations, similar to scATOMIC. Scmap-cell^15^ classification was performed using default parameters. To enable SingleR processing the large reference dataset we applied its pseudobulk implementation by setting ‘aggr.ref’ to TRUE^13^. Otherwise, default parameters were used. To compare scATOMIC’s final cancer cell prediction with the CNV inference approach of detecting malignant cells we used CopyKAT^17^ with its default settings (Fig. 3). Both aneuploid and diploid cells from the external validation biopsies were included in this analysis. Agreement rate was defined as the simple percentage agreement using the agree function from the irr^47^ R package. Cells which received an NA ploidy annotation were omitted from the calculation.

### Breast cancer subclassification

The model extending breast cancer into its molecular subtypes was trained using a combination of two datasets. The first included the breast cancer cell lines data within the core training dataset (Supplementary Table 1,2), where the assigned molecular subtypes were based on annotations found in the Cancer Cell Line Encyclopedia DepMap portal^48^ and the second, primary tumour data from *Wu et al*^49^ with immunohistochemistry information (Supplementary Table 4). We used the clinical molecular subtypes of HER2+, ER+ and triple negative as classes. There was not sufficient data available to train and test HER2+/ER+ and HER2-/ER+ as separated classes. We evaluated this extension’s performance on an external set of 40 tumours from *Pal et al*^35^ (Supplementary Table 5). One tumour containing fewer than 100 cancer cells was omitted from this analysis. The ground truth of ER and HER2 status in the testing set was received by correspondence with the authors.

### Visualizing inferred CNVs

To visualize the inferred CNV profile in the ER-low tumour we used the ‘inferCNV’^50^ package with the cutoff variable set to 0.1 and all other variables set to default. We defined normal reference cells as cells annotated by scATOMIC as blood or stromal cells.

### Applying scATOMIC to metastatic tumours

scATOMIC was applied to predict the tumour of origin in 62 metastatic tumours. Individual samples from are described in detail in Supplementary Table 6. We defined the scATOMIC tumour of origin prediction by taking the cancer type called in the majority of cancer cells in the sample.

## Acknowledgements

This work was supported by a grant from The Banting Research Foundation and Investigator Awards received from the Ontario Institute for Cancer Research with funds from the province of Ontario to SA and PA. INM obtained funds from the Ontario Graduate Scholarship Program - University of Toronto. We thank Salman Basrai for his assistance checking the validity and accuracy of the scATOMIC code, as well as on his comments and suggestions concerning the package documentation.

## Author Contributions

All authors discussed the results, wrote and commented on the manuscript. INM developed the concept, performed data analysis, developed the tool and wrote the code. DS contributed to statistical analysis and model development. PA and SA conceived the idea, contributed to data analysis, led, and supervised all aspects of the study.

## Code availability

scATOMIC’s associated code and user manual are available at https://github.com/abelson-lab/scATOMIC

## Extended Data Figures

**Supplementary Fig. 1.**
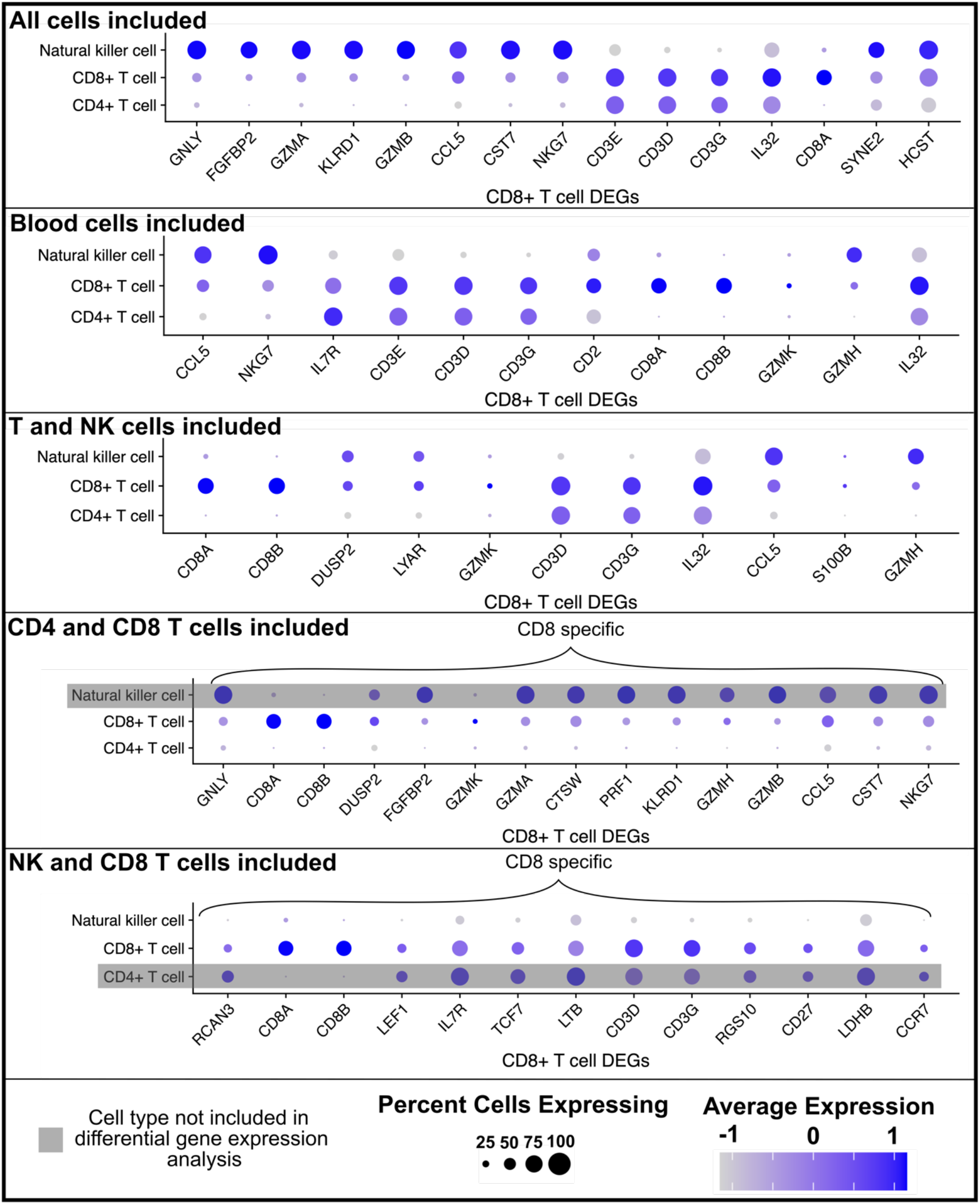
Expression of CD8+ T cell marker genes in related cell populations. The expression of the top CD8+ T cell markers derived from differential gene expression analysis are shown. Features that distinguish CD8+ T cells from CD4+ T cells are highly expressed in natural killer cells and vice versa.

**Supplementary Fig. 2.**
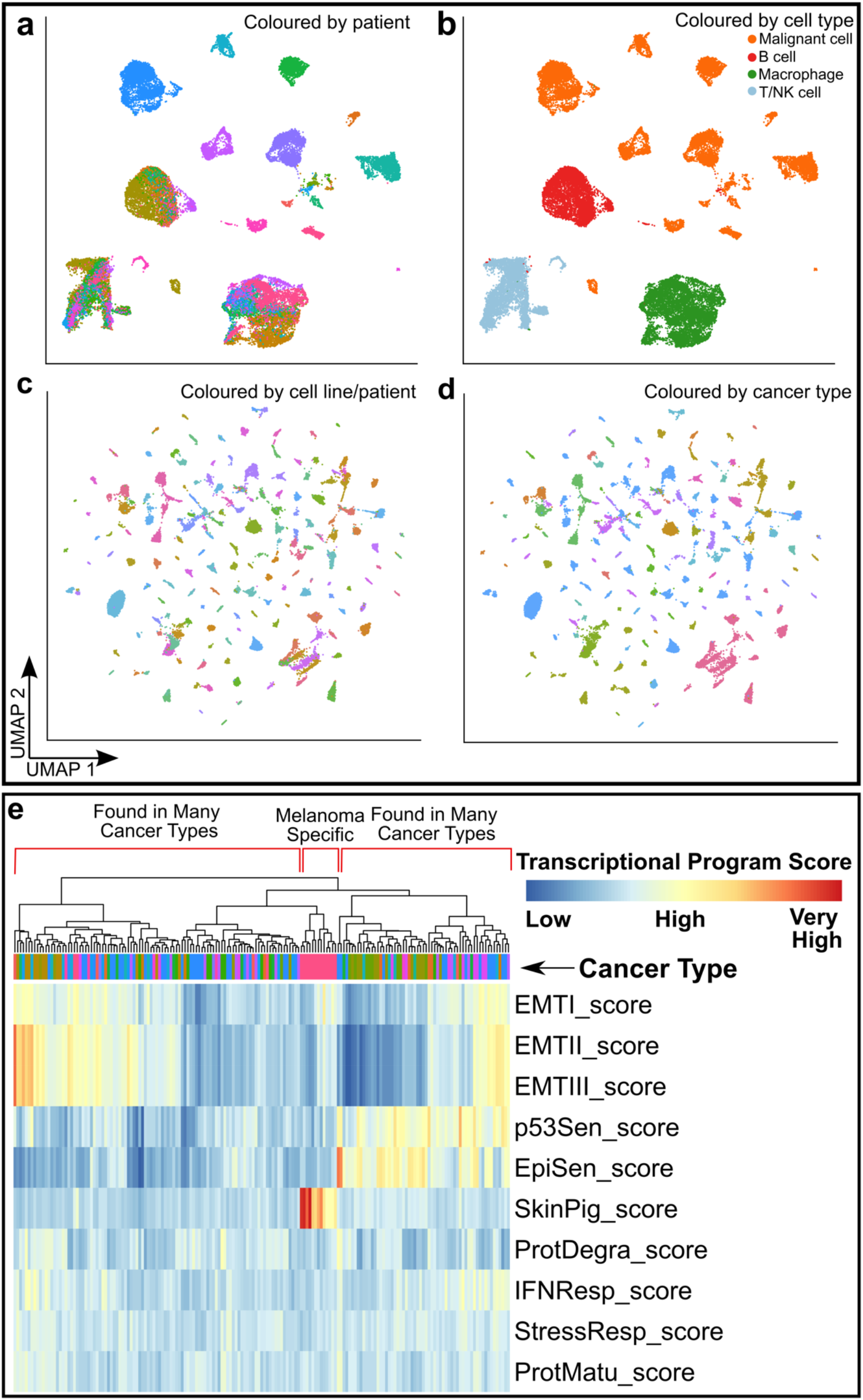
Complexity associated with interpatient tumour cell heterogeneity. **a**,**b**, Unsupervised clustering of 17 lung cancer biopsies^51^. Dots represent single cells coloured by **a**, patient ID, and **b**, the reported cell type. **c**,**d**, Unsupervised clustering of 175 cancer cell lines’ scRNA-seq data. Dots represent single cells and are coloured by **c**, cell line, and **d**, cancer type. The transcriptome of each patient derived malignant cell population/cell line is unique, thus whole transcriptomes don’t cluster cancers by type, while immune cells from different patients co-cluster. **e**, Hierarchical clustering based on the expression of established cancer associated transcriptional programs^18^. Transcriptional programs reflecting diverse biological processes in cancer are largely independent of the cancer type. Programs’ enrichment scores were obtained directly from the manuscript.

**Supplementary Fig. 3.**
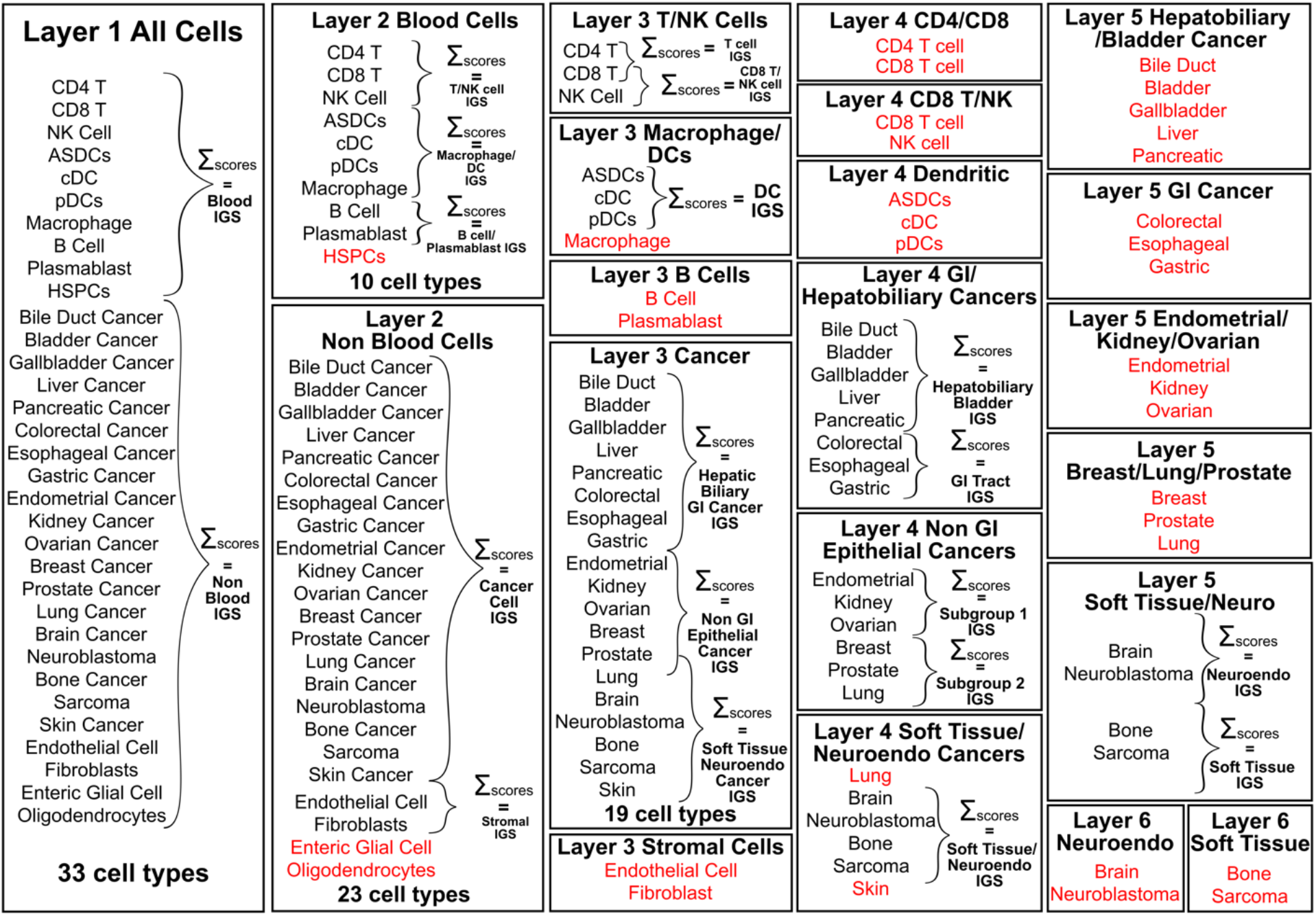
Detailed overview of classification branches and intermediate-group scores within the core scATOMIC algorithm. Random forests classification models representing different parental nodes (n=21) are used to derive scores for terminal cell types. Broad cell type intermediate-group scores are calculated by taking the sum of random forest scores for each terminal cell type indicated. Red text represents cell classes that can reach terminal classification in each model.

**Supplementary Fig. 4.**
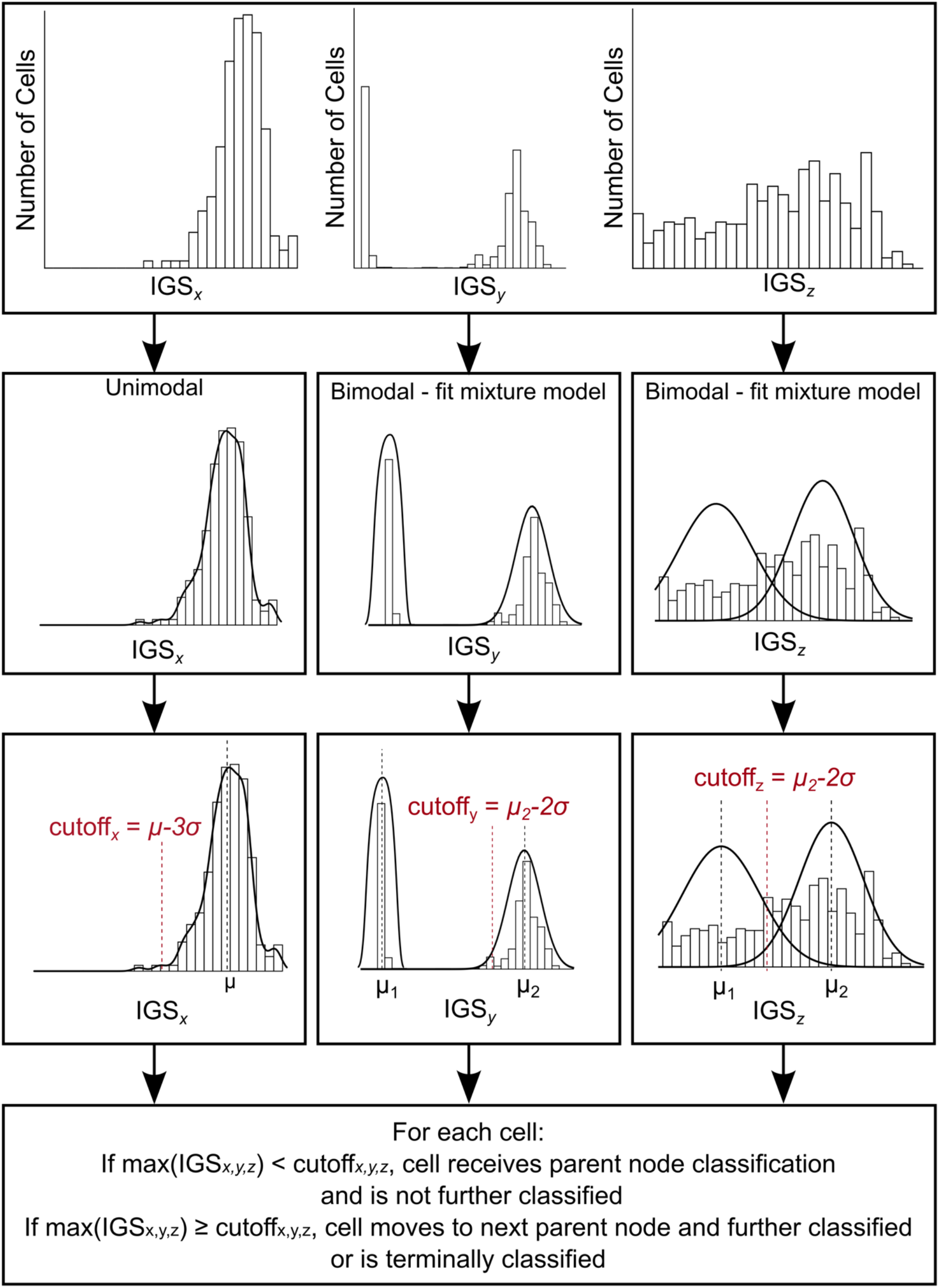
Setting automatic IGS cut-offs at each classification branch. IGS cut-offs are determined based on the distribution of scores derive from all the cells being queried in each particular classification task. The modality of IGS distributions is classified as either unimodal or bimodal. For unimodal distributions the cut-off to associate one cell with a cell class is set to be three standard deviations (σ) from the mean (µ). For bimodal distributions scATOMIC fits a mixture model and estimates parameters using the expectation maximization algorithm. The cut-off is set to 2 standard deviations from the highest estimated mean (µ_2_). IGSx, IGSy, IGSz represent possible distributions of scores obtained from different models. For example, unimodal distribution can be seen when highly purified cell population is being interrogated by a particular model. 2-mode bimodal distribution is typical for the first classification task (i.e., Blood or Non-blood) of cancer TMEs.

**Supplementary Fig. 5.**
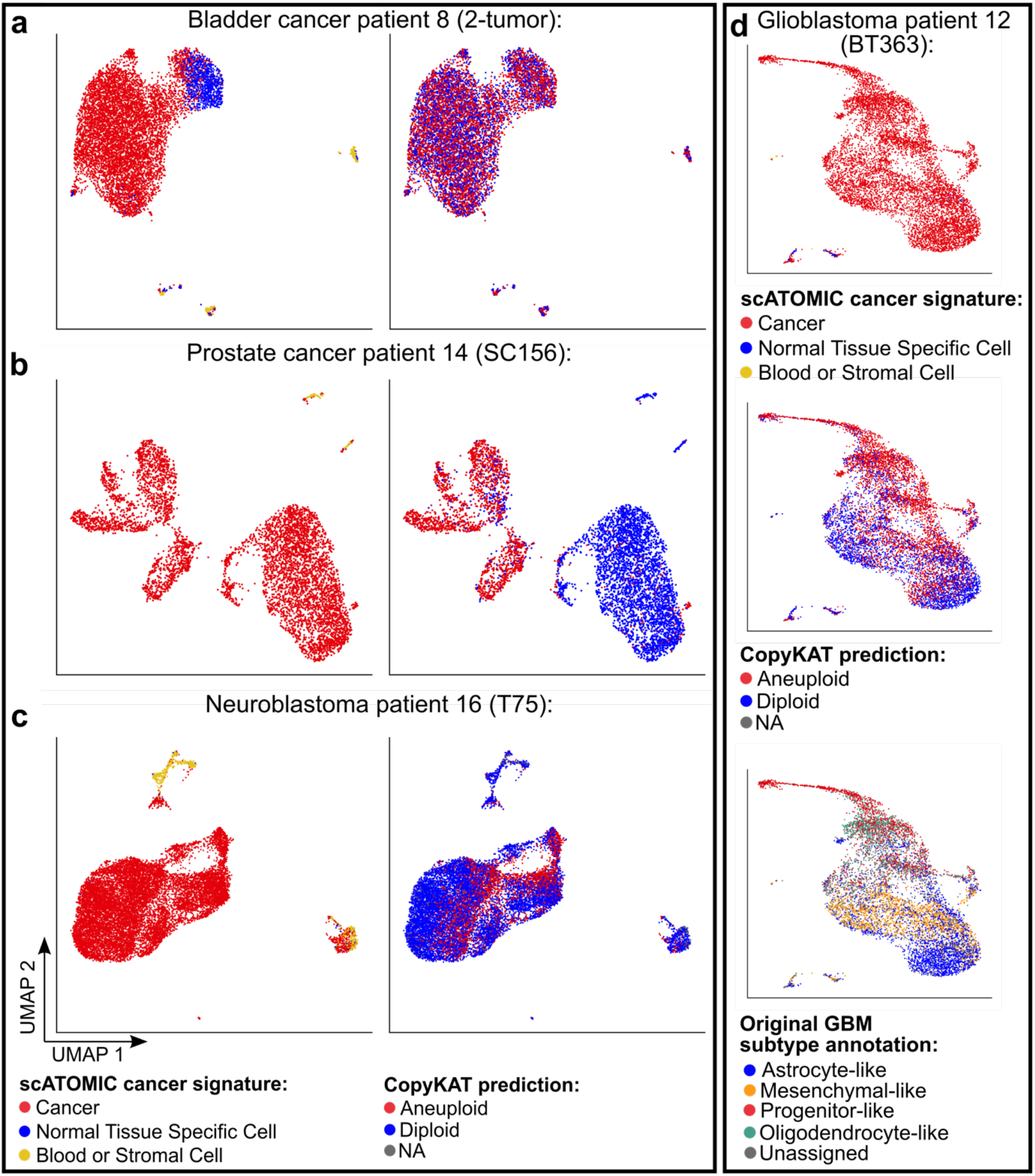
Examples of discordant scATOMIC malignant status and CNV-based inferred ploidy. Cells are coloured according to scATOMIC malignant annotation and CopyKAT inferred CNV status. **a**, A bladder tumour reported to have low CNV burden^4^. **b**, A prostate tumour specimen with few reference cells. The specimen reported to have 80% cancer cellularity^31^. **c**, A neuroblastoma tumour with an un-patterned prediction of aneuploid cells suggesting weak CNV profile intensities above the reference^52^. **d**, A glioblastoma tumour with multiple tumour cell populations^53^.

**Supplementary Fig. 6.**
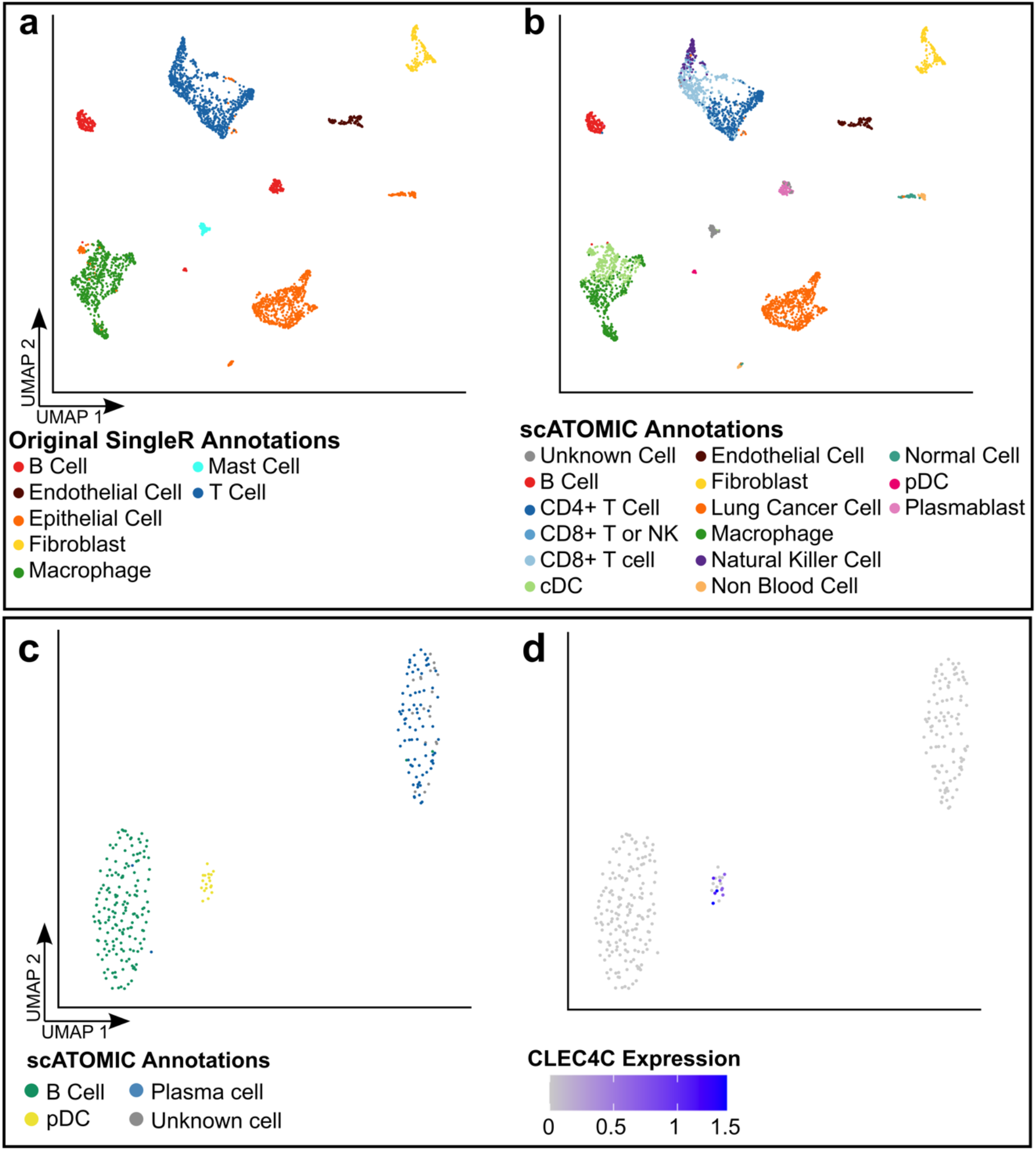
Increasing the cellular resolution with scATOMIC’s core predictions. scATOMIC was applied to a published lung cancer biopsy dataset^28^. Cells were clustered and visualized using the standard Seurat workflow and colored by **a**, the original published annotations^28^ and **b**, scATOMIC predictions. **c**, Cells that were originally annotated as B cells are illustrated on two UMAP dimensions. Single cells are labelled by scATOMIC annotations. scATOMIC separated B cells into B cells, Plasma Cells, and pDCs. **d**, Single cells are labelled by the expression level of the pDCs marker *CLEC4C*. Expression values represent normalized read counts.

**Supplementary Fig. 7.**
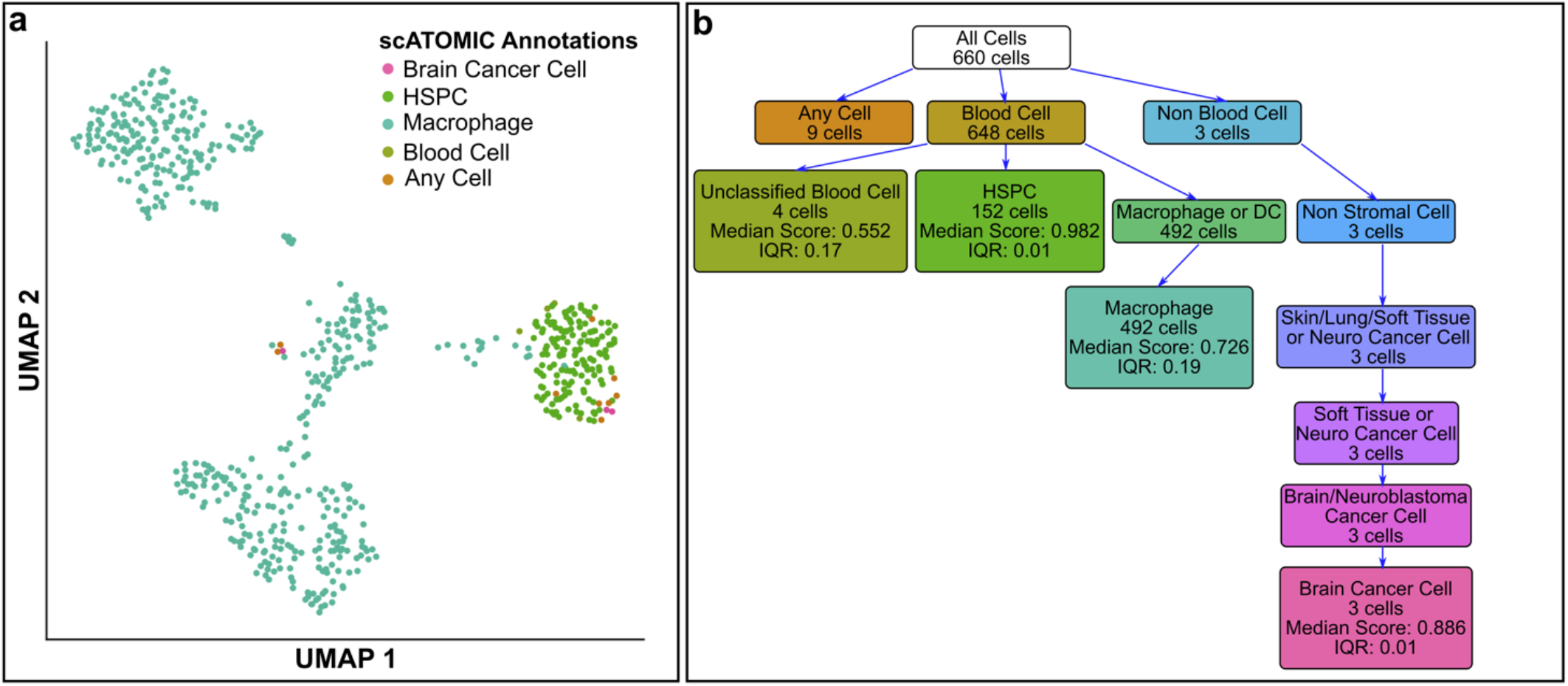
scATOMIC identifies HSPCs in glioblastoma. We applied scATOMIC to a dataset of CD45+/CD34+ bead enriched cells from primary glioblastoma tissue. **a**, UMAP illustration of scATOMIC predictions. **b**, scATOMIC correctly identified HSPCs and separated those from other cell types, including macrophages and 3 brain cancer cells, thus recapitulating the original cancer type reported by the authors^33^.

**Supplementary Fig. 8.**
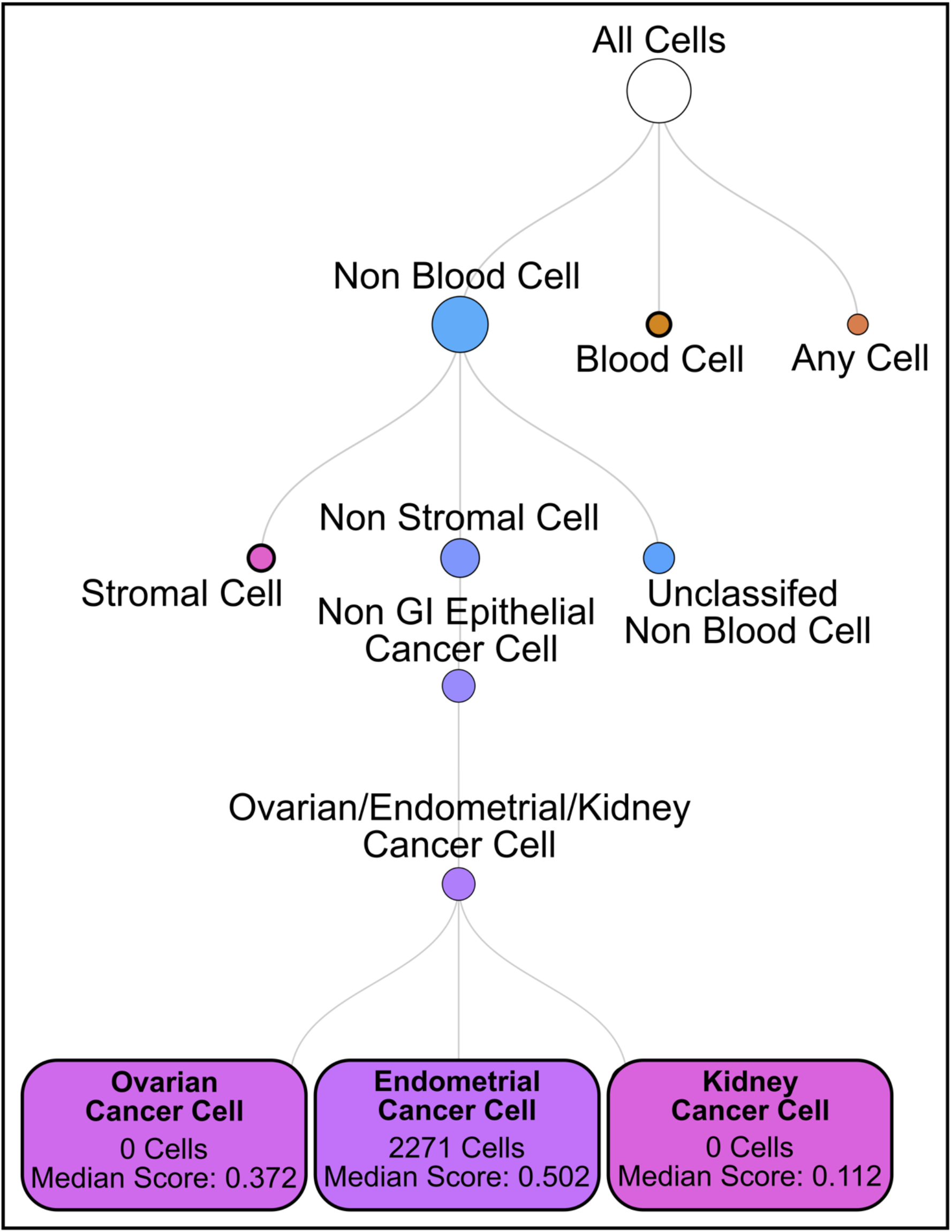
Example for a misclassification of an ovarian cancer specimen. The classification hierarchy for cancer cells is shown with increasing resolution down the tree. Median classification scores, (that is, the proportion of trees voting for each class), are shown for kidney, endometrial, and ovarian cancer cells in the final branch.

## Description of Supplementary Tables

**Supplementary Table 1** – Reference datasets used to train scATOMIC’s core cell class models. For each cell type in the reference the dataset source, number of cells, and mean F1 score in k-fold cross validation is shown.

**Supplementary Table 2** – List of the cell lines from *Kinker, G*.*S. et al (2020)* used in the reference dataset. For each cell line, the number of cells, cancer type, and cancer subtype is shown.

**Supplementary Table 3** – Pan cancer external validation datasets. For each sample the source of data, condition, cell types, and number of cells is shown. F1 scores for scATOMIC, scmap-cell, SingleR, SingleCellNet, and CHETAH are shown for each cell type from each sample.

**Supplementary Table 4** – Training set samples for breast cancer subclassification. Immunohistochemical subtype and the number of cells is shown for each sample.

**Supplementary Table 5** –Validation breast cancer samples from *Pal, et al (2021)*. For each sample, the reported subtype, predicted subtype, and cell composition is shown.

**Supplementary Table 6** – List of annotated metastatic samples used for tumour tissue of origin prediction.

**Supplementary Table 7** – List of features being used by scATOMIC for cell classification. Features are ordered to indicate genes that differentiate one cell type from all the other cells represented in the same layer.

